# Field competitiveness of *Aedes albopictus* [Diptera: Culicidae] irradiated males in pilot Sterile Insect Technique trials in Northern Italy

**DOI:** 10.1101/2020.09.25.313544

**Authors:** Romeo Bellini, Marco Carrieri, Fabrizio Balestrino, Arianna Puggioli, Marco Malfacini, Jeremy Bouyer

## Abstract

Vector-borne diseases account for 17% of infectious diseases, leading to more than one million deaths each year. Mosquitoes are responsible for 90% of the casualties and alternative control methods to insecticides are urgently needed, especially against *Aedes* vectors. *Aedes albopictus* is a particularly important species, causing major public health problems because it is a vector of several arboviruses and has a strong invasive behaviour. Various genetic control methods have been proposed to be integrated into the management strategies of *Aedes* species, among which the sterile insect technique (SIT), which proved efficient against various insect pests and vectors. However, the ability of released irradiated sterile male mosquitoes to compete with their wild counterparts and induce sterility in wild females, which is critical to the success of this strategy, remained poorly defined. Here, we assessed the field competitiveness of *Ae. albopictus* irradiated male using data from six release trials implemented in Northern Italy for three years. Sterile males were capable of inducing a good level of sterility in the wild female population, however with high variability in time and space. The field competitiveness of the released males was strongly negatively correlated with the ratio of sterile to wild males. This should be taken into consideration when designing future programmes to suppress field populations of *Aedes* mosquitoes.

## Introduction

In the recent document on the global vector control response 2017-2030, the World Health Organization (WHO) pointed out the urgent need for alternative mosquito control methods, particularly against *Aedes* vectors (WHO 2017). The main reason behind this need is that the integrated control of these vectors as proposed until now is challenging in many countries and in different climatic conditions without guarantying satisfactory levels of population reduction (Reiter 2016). Moreover, the EU Biocide directive is progressively restricting many insecticides in Europe, reducing *de facto* the vector control options and increasing the probability for the development of resistance against the remaining insecticides (Grigoraki et al. 2017, Pichler et al. 2018). New techniques to control urban mosquitoes are therefore under development and field evaluation, such as genetic control strategies targeting the reproductive capacity of disease transmitting mosquitoes (Flores and O’Neill 2018, McGraw and O’Neill 2013). Amongst those, the sterile insect technique (SIT) shows great promise but requires the production of large quantities of mosquitoes in adequate facilities. The male mosquitoes are sterilized using ionizing radiation that generates random dominant lethal mutations in the germinal cells. When released in large numbers to outcompete their wild counterparts, sterile males will mate with wild females and sterility is induced due to embryonic arrest. There will be no offspring, thus reducing the population growth rate in the next generation (Dyck et al. 2005). This technique has been used very successfully against various agricultural pests (Wyss 2006, Enkerlin et al. 2015) and vectors (Dicko et al. 2014, Vreysen et al. 2000), and has been under development for several years against disease transmitting mosquitoes with major progresses reported recently (Lees et al. 2020). Thirty four pilot SIT trials against mosquitoes are reported as presently ongoing worldwide (Bouyer et al. 2020a).

*Aedes albopictus* is a major public health concern as it is a very good vector of several arboviruses such as dengue, chikungunya and Zika (Mitchell 1995, Wong et al. 2013). The potential epidemiologic consequences of adaptation to *Ae. albopictus* mosquitoes are well-documented for chikungunya virus, for which some mutations of the virus increased vector competence (Tsetsarkin et al. 2007). The epidemiological risk related to *Ae. albopictus* has become a reality in temperate Europe, where a number of disease outbreaks have occurred in recent years (Gossner et al. 2018). Feasibility studies on the use of the SIT for targeting the invasive *Ae. albopictus* mosquito were started in Italy in 2000 (Bellini et al. 2007), including several release field trials with irradiated males that produced large data sets (Bellini et al. 2013a).

## Materials and Methods

### Study area

Between 2008-2018, eight release trials with irradiated *Aedes albopictus* males were carried out in several localities in Northern Italy to test their efficacy in population suppression. The selected urban localities were representative of the urban conditions in Northern Italy. They were also well-isolated from other urban areas by agricultural land and with size adequate to fit the local sterile male production capacity.

The main characteristics of these trials are presented in the Tab. 1.

**Tab. 1.** Main descriptive data from the eight SIT field release trials on *Aedes albopictus*

| Locality and release year | Sterile male strain | *Larval Diet | Dose GY | Stage released | Release period (from-to) | N. release station/ha | N. releases | Release area (ha) |
| --- | --- | --- | --- | --- | --- | --- | --- | --- |
| Boschi 2008 | Rimini F26 | CAA | 30/40 | pupae | Apr 29-Oct 02 | 0.62 | 29 | 16 |
| Budrio 2008 | Pinerolo F25 | CAA | 40 | pupae | Apr 30-Aug 08 | 0.70 | 18 | 17 |
| Boschi 2009 | Rimini F35 | CAA | 30 | pupae | May 14-Sep 10 | 0.62 | 14 | 16 |
| Caselline 2009 | Pinerolo F35 | CAA | 30 | pupae | May 21-Sep 25 | 0.61 | 19 | 18 |
| Boschi 2010 | Rimini F41 | IAEA | 30 | pupae | May 14-Sep 14 | 0.25 | 19 | 16 |
| Gherghenzano 2012 | RER F7 | IAEA-BY | 30 | pupae & adults | May 15-Sept 25 | 1.00 | 20 | 10 |
| Caselle 2018 | Rimini F68-70 | IAEA-BY | 35 | adults | June 15-Aug 31 | 1.00 | 14 | 16 |
| Guisa Pepoli 2018 | Rimini F68-70 | IAEA-BY | 35 | adults | June 15-Aug 31 | 1.00 | 12 | 7 |

\*CAA diet—larval diet consisting of 80% Friskies dry adult cat food, 14% brewer’s yeast, and 6% Tetramin fish food. IAEA diet—larval diet consisting of 50% bovine liver powder, 50% tuna meal, and 0.4% w/v of Vitamin Mix. IAEA-BY diet— larval diet consisting of 50% bovine liver powder, 36% tuna meal, 14% brewer’s yeast and 0.2% w/v of Vitamin Mix.

### Mass rearing and strains

Different mosquito strains and generations were used in the pilot trials, but in all cases, colonies were started from eggs collected in Northern Italy (Tab. 1). The mosquitoes were reared under standard holding conditions: 27±2°C, 85% RH, and a photoperiod of 14:10 (L:D) h, as described in Bellini et al. (2013a).

### Male sorting, sterilization and release protocol

Separation of the sexes was carried out on the pupae in water by means of metal sieves with a mesh size of 1,400 μm. The male insects were sterilized by exposing 24-36 h old male pupae to 30-40 Gy in an IBL 437 Cobalt irradiator (CIS Bio International, Bagnols-sur-Ceze, France) (Bellini et al. 2013a). The male pupae were released 1-2 h after irradiation into plastic containers positioned on the ground in permanent shaded sites. In Gherghenzano, each batch of irradiated pupae was divided into two and one half was released as pupae; the other half was taken to the laboratory and emerged adults were released three days later. In Caselle and Guisa Pepoli, adults were ground released in fixed stations distanced about 100 m from each other.

### Field data collection

In the period from 2008-2012, ovitraps CAA7 consisting of black plastic pots of 400 ml (upper diameter 8 cm, lower diameter 6 cm) holding about 250 ml dechlorinated water and one 12.5 x 2.5 cm Masonite strip were used for egg oviposition. In 2018, ovitraps CAA14GR consisting of black plastic cylindrical pots of 1,400 ml holding 800 ml of dechlorinated water, with a grill screen to prevent access to the animals and a strip of masonite (15×2.5 cm) as egg deposition substrate. The traps were deployed in the release as well as in the control areas and checked weekly (Tab. 2). Eggs were counted under a stereomicroscope and hatched using standard procedures (Bellini et al. 2007) to check for their fertility rate (% egg hatch).

**Tab. 2.**
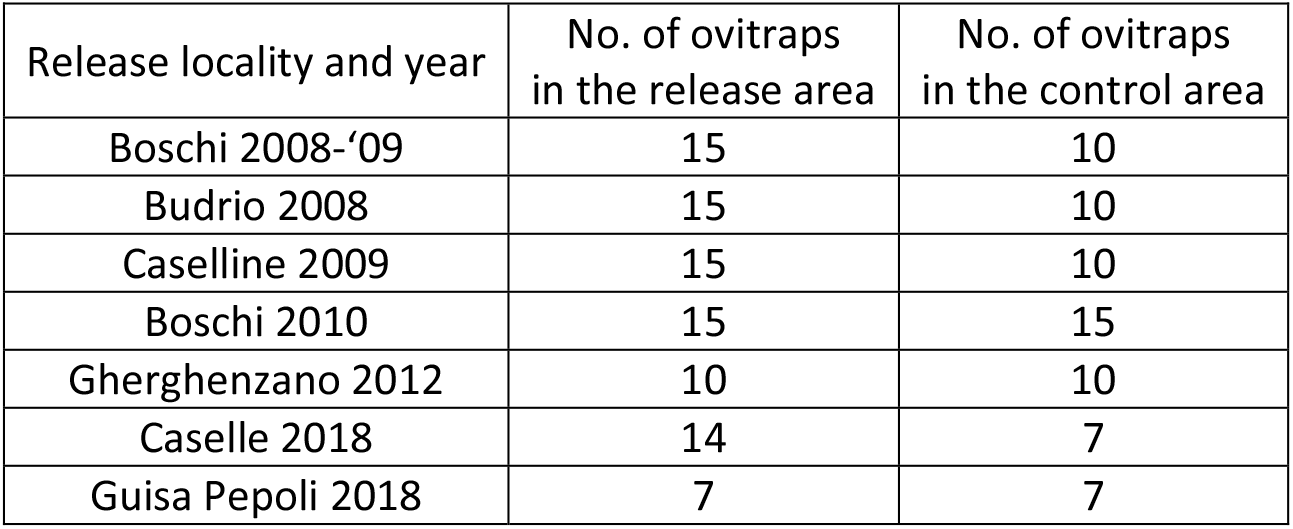
Number of ovitraps used in the release trials

| Release locality and year | No. of ovitraps in the release area | No. of ovitraps in the control area |
| --- | --- | --- |
| Boschi 2008-'09 | 15 | 10 |
| Budrio 2008 | 15 | 10 |
| Caselline 2009 | 15 | 10 |
| Boschi 2010 | 15 | 15 |
| Gherghenzano 2012 | 10 | 10 |
| Caselle 2018 | 14 | 7 |
| Guisa Pepoli 2018 | 7 | 7 |

### Statistical analysis

The percentage of egg-induced sterility (S) was calculated in relation to the number of hatched eggs in the control area using Abbott’s equation:

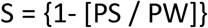

where PS and PW are the percentages of hatched egg in the release and in the control area, respectively.

The percentage of decrease of egg collection (D) in the release area in relation to the control area was calculated by the following equation:

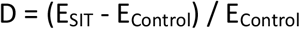

where E_SIT_ and E_Control_ are the mean number of eggs per ovitrap per week in the release and in the control area, respectively (see also supplementary data).

The t-test for dependent samples was performed to compare the results obtained by the methods of estimation of the sterile/wild ratio with the observed data.

The linear regression analysis was used to determine the relationship between the adult caught by HLC and the weekly number of eggs collected by ovitraps.

### Estimation of wild and sterile male densities

The estimation of wild and sterile male densities required to calculate the field competitiveness was conducted with three methods.

#### Method 1

Wild male population density was estimated based on the relation between the mean number of eggs collected by ovitraps CAA14GR and the male density estimated in three urbanized areas in Bologna city (Italy) in 2011. In the three areas (total 841 hectares), all breeding sites present in 610 premises (equaling 9.82% of the total premises present in the area) were sampled and 15 ovitraps were activated. Each urban area was divided into 13-16 zones and 15 premises per zone were randomly selected for thorough inspection. During the inspections, all *Ae. albopictus* L4 larvae and pupae were collected in all breeding sites, counted and classified. The male density was estimated based on the number of sampled L4 larvae and pupae using the model developed by Vallorani et al. (2013) and considering a sex ratio (F:M) of 1.08 (0.6 SD) which was calculated in a control area during a mark-release-recapture (MRR) study conducted in 2018 (supplementary data). The relationship between the wild male/ha (Mw) and the number of eggs/ovitrap/week (E_CAA_) is given by:

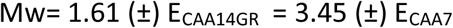

where E_CAA14GR_ and E_CAA7_ are the number of eggs per model ovitrap CAA14GR (biweekly collection) and CAA7 (weekly collection), respectively (the rate E_CAA14GR_/ E_CAA7_ = 2.14 (±0.31 SD)) (Carrieri et al. 2017).

The daily sterile male population density was estimated, taking into account the number of sterile males released and their daily survival rate (SR) that was estimated by the mean daily relative humidity (RH) as described in Bellini et al. (2010) using the equation:

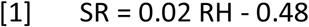

This equation is considered valid in the RH range 48-72.5 %; above RH=72.5%, SR is assumed to be equal to 97%, and below RH=48%, SR is assumed to be equal to 52% (data set supporting this approach is available in Supplementary data).

To estimate more precisely the S / W males ratio, the number of sterile males that survived in the previous release (*Mss*) were considered by adding their number (estimated by the equation 1) to the wild males (*Mw*) in order to be comparable to the values observed in MRR studies (marked males/wild males + sterile males from the previous week).

#### Method 2

*Ae. albopictus* wild male and female population densities were estimated by human landing collection (HLC) using manual aspirators in the release and control areas in MRR studies. In 2018 two sampling sessions were conducted in parallel in the release (Guisa Pepoli and Caselle) and in the control areas (Bolognina). The male to female ratios (M / F) in the release and in the control areas were calculated and compared to determine the sterile to wild males ratio in the release area for each egg monitoring week using the equation:

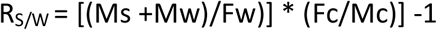

where R_S_/_W_ is the ratio of sterile/wild males; Ms, Mw and Fw are the numbers of sterile males, wild males and wild females, respectively, collected in the release area; Fc and Mc are the numbers of females and males, respectively, collected in the control area.

Method 2 has also been corrected to compare it with the rate observed in MRR studies; the rate Rs/w calculated were multiplied by (1-Mss)/Mw.

#### Method 3

The ratio of sterile (colored) to wild males were calculated in MRR studies to verify the outcome of methods 1 and 2 (Supplementary data).

Two MRR sessions were undertaken in 2018: the first in the period July 06-13 and the second between August 03-10 (supplementary data). Marked sterile males were released in the SIT treatment areas only (Guisa and Caselle) while the recapture sessions were performed in parallel either in the release or in the control localities (Bolognina).

Twenty-four sampling stations were randomly selected in each locality. The field sample collections, using a manual aspirator for 5 minutes in each sampling station, were conducted daily from 5:00 PM to 7:00 PM, starting from the first day after release and continued for seven consecutive days.

### Field competitiveness of irradiated males

Field competitiveness was estimated through the weekly capacity to induce sterility (CIS) index, a simplified version of Fried’s competitiveness index (Bellini et al. 2013b). CIS was calculated using the following equation:

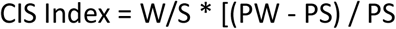

where W and S are the numbers of wild and sterile males, respectively; PW is the percentage egg hatch in the control area and PS is the percentage egg hatch in the release area.

Dependent-samples T-test was used to compare the two methods for estimating the sterile to wild male ratios. For describing the central tendency of the sterile/wild ratio, we also used the median because it was less affected by outliers.

Method 1 was used to estimate the sterile to wild males ratio for the trials carried out in all periods (2009-2018).

## Results

Results showed that sterile males released weekly at the dose range of 900-1500 males/ha/week induced sterility levels from 15 to 70% in the local egg population. When induced egg sterility reached 70%, a similar reduction was found in egg numbers in the ovitraps.

The field data were analyzed to estimate the field competitiveness of the released *Ae. albopictus* sterile males.

The sterile to wild male ratios observed during the field trials using the two methods applied are shown in Table 3.

**Tab. 3.** Estimation of the weekly sterile to wild male ratios using the two applied methods and data observed in MRR. Field data (mean ± SD) were collected in Guisa and Caselle (SIT area) and Bolognina (control area) in 2018

| Ratio<br>Sterile/wild | Mean | SD | N | Diff. | S.D.<br>Diff. | T-test for dependent samples |  |  | Product Moment<br>Correlation |  |
| --- | --- | --- | --- | --- | --- | --- | --- | --- | --- | --- |
|  |  |  |  |  |  | t | Df | p | R | P |
| **Observed** | 0.65 | 0.82 |  |  |  |  |  |  |  |  |
| **Method 1** | 0.69 | 0.53 | 24 | -0.04 | 0.45 | -0.42 | 23 | 0.68 | 0.87 | <0.001 |
| **Method 2** | 1.07 | 1.52 | 24 | -0.42 | 1.14 | -1.79 | 23 | 0.09 | 0.68 | <0.001 |

The mean values of the weekly sterile to wild male ratios in 2018 were similar for the two methods used and no significant difference was observed when the two methods were compared with observed MRR data. The sterile to wild male ratio S/W calculated with method 1 was thereafter used to assess the competitiveness of the sterile males in all the other pilot trials. The mean CIS index strongly varied in space and time ranging from 0.02 to 0.37, which indicates that the sterile males were 3 to 100 times less competitive than the wild males (Table 4).

**Tab. 4.** Data from eight sterile male release trials carried out in Northern Italy. The capacity to induce sterility (CIS) was estimated weekly.

| Area_year | No. sterile males released/ha |  | D (% egg reduction) |  | S (% induced egg sterility) |  | Weekly CIS index |  | Rate sterile/wild |  |
| --- | --- | --- | --- | --- | --- | --- | --- | --- | --- | --- |
|  | Mean | SD | Mean | SD | Mean | SD | Mean | SD | Mean | SD |
| Boschi_2008 | 1475.1 | 1005.2 | -47.38 | 25.20 | 56.02 | 28.42 | 0.29 | 0.33 | 51.55 | 89.92 |
| Boschi_2009 | 592.0 | 295.5 | -73.03 | 12.39 | 64.75 | 18.24 | 0.67 | 1.02 | 35.36 | 109.14 |
| Boschi_2010 | 857.8 | 236.3 | 6.84 | 61.24 | 16.00 | 31.60 | 0.11 | 0.15 | 5.69 | 5.78 |
| Budrio_2008 | 1632.0 | 959.4 | 20.04 | 81.43 | 48.08 | 22.56 | 0.67 | 1.18 | 72.92 | 195.06 |
| Casellina_2009 | 2800.3 | 9288.0 |  |  | 48.65 | 25.51 | 0.89 | 1.00 | 37.43 | 100.53 |
| Gherghenzano_'12 | 10659.9 | 6842.8 |  |  | 50.98 | 19.18 | 0.04 | 0.04 | 105.17 | 135.74 |
| Guisa_2018 | 783.1 | 545.1 | -52.61 | 17.77 | 22.23 | 23.44 | 0.19 | 0.43 | 4.27 | 3.95 |
| Caselle_2018 | 226.2 | 97.0 | -29.28 | 23.69 | 5.02 | 14.46 | 0.16 | 0.32 | 0.72 | 0.40 |
| All Grps | 2870.8 | 5732.9 | -33.33 | 52.92 | 42.42 | 29.42 | 0.39 | 0.75 | 44.82 | 111.02 |

\* The percentage decrease of egg collection (D) was calculated using the following equation: $D = (E_{Control} - E_{SIT}) / E_{Control}$ where $E_{SIT}$ and $E_{Control}$ are the mean number of eggs per ovitrap per week in the release and in the control area, respectively. The percentage of egg sterility in SIT and control areas was estimated by calculating egg hatch using standard procedures, as reported in Bellini et al. (2013b).

The capacity to induce sterility was highly variable in the eight field pilot trials analysed, with a mean value of CIS 0.39 (SD±0.75) and a median value of 0.13 (0.02-0.39 Q_25-75_). A strong temporal variability was observed when plotting the CIS index against the seasonal period, with lower CIS values observed at the beginning and at the end of the season, when the wild population density is usually lower (Albieri et al. 2010, Carrieri et al. 2011a, 2011b) and therefore the ratio of sterile to wild males tends to be higher (Fig. 1).

**Fig. 1.**
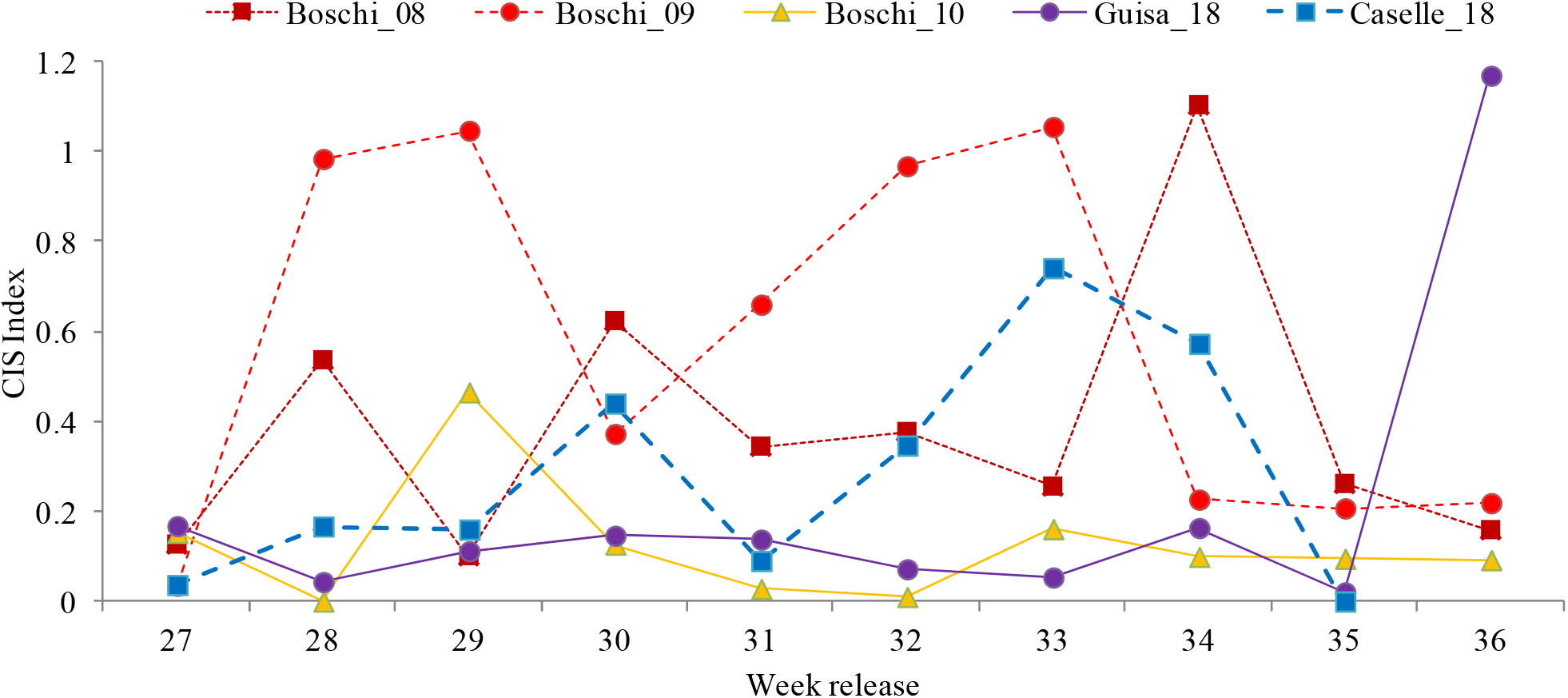
Seasonal dynamics of the weekly CIS index during field pilot trials conducted in Boschi (2008, 2009 and 2010) and in Guisa and Caselle in 2018.

These low values of the CIS index observed at the beginning and at the end of the season may indicate that the released males were less prone to disperse and mate due to lower temperatures during these periods in comparison with the higher temperatures in the middle of the summer, thus reducing the actual sterile to wild males ratio with distance from the release points. Competitiveness of the sterile males might also be density-dependent as indicated by the strong negative correlation between CIS values and ratios of sterile/wild males as expressed by the equation: Ln R_S/W_ = 0.39 – 0.73 Ln (CIS index) (R^2^ = 0.62, F_(1,113)_= 185.26, p <0.0001).

**Fig. 2.**
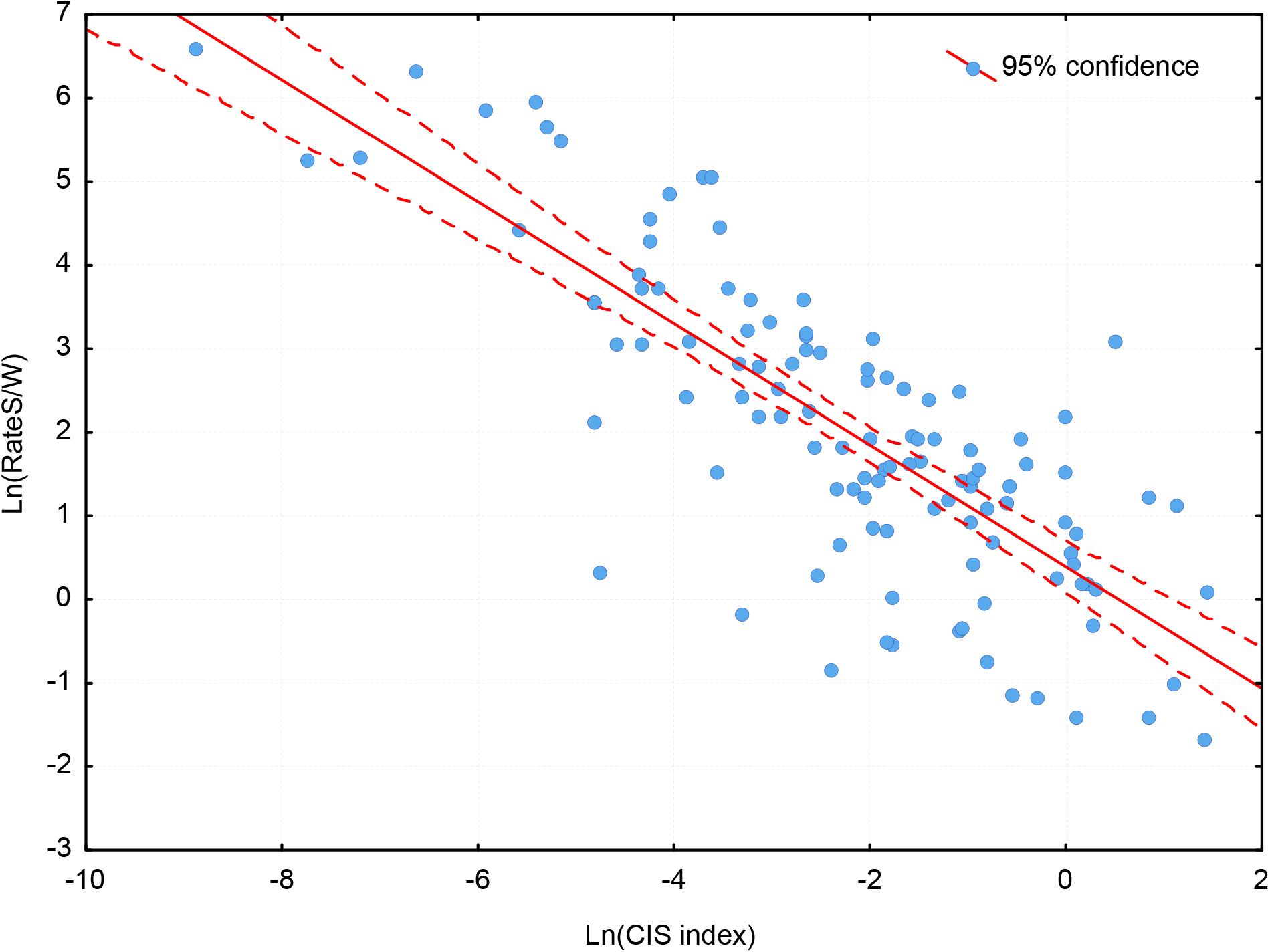
Linear regression between Ln (S/W) males and Ln (CIS values). This analysis included all weekly results and all sites.

## Discussion

The competitiveness of the sterile males used in the field trials analysed in the present study was tested in previous large semi-field cages showing a CIS index of 0.96 ± 0.62 and 0.71 ± 0.36 at 30 and 40 Gy, respectively (Bellini et al. 2013b), indicating the good quality of the sterile males released.

A competitiveness index of at least 0.2, measured in semi-field cages with a ratio between sterile to fertile males of 1: 1, is considered acceptable in Tephritid flies SIT programs (FAO/IAEA/USDA, 2019).

When moving from semi-field setting to the field, a reduction in the competitiveness index is expected for several reasons. These include: the pulsing release of sterile males (mainly weekly releases in this case) while wild males emerge every day; the daily mortality of sterile males causing their density to decline during the intra-release time; the capillary unknown distribution of breeding sites matched with the point site release of sterile males; the obstacles to the dispersal of sterile males caused by artificial barriers in urban areas; and the immigration of already mated females from outside the SIT pilot area.

The median CIS index obtained in the eight SIT field trials conducted in Northern Italy in the period 2008-2018 was 0.12 (range 0.03 – 0.38), which is considered good when compared with competitiveness indexes observed in other genetic control pilot field trials.

In pilot trials conducted in Brazil and the Cayman Islands by releasing the transgenic *Ae. aegypti* strain OX513A, field competitiveness values of 0.031 (95% CI: 0.025-0.036) and 0.059 (95% CI: 0.011 – 0.210), respectively, were obtained (Harris et al. 2011, Carvalho et al. 2015). The field competitiveness of *Ae. albopictus* irradiated mosquitoes assessed in our study was thus overall much higher than that of *Ae. aegypti* transgenic strains.

The large variability observed in the CIS index among releases and among weeks of the same trial is difficult to explain and may be attributable to the impact of climatic condition.

We hypothesized that part of the wild females might be unattainable by the sterile males whatever the ratio imposed because they are located in cryptic habitats difficult for the sterile males to access. This might be particularly relevant in urban areas because of the numerous artificial obstacles such as perimetric hedges and walls. It will probably be possible to reduce this gap by air releasing sterile males in order to achieve a more homogeneous covering of the target area (Bouyer et al. 2020b).

In natural environments, the field competitiveness of *Glossina palpalis gambiensis* sterile males was estimated 0.07 in Burkina Faso (Sow et al. 2012), while it resulted 0.14 (SD 0.10) and 0.76 (SD 0.11) for two different strains in Senegal (Bassène et al. 2017).

In the case of the New World screwworm (*Cochliomyia hominivorax*), field competitiveness was estimated at 0.29–0.43 at smaller scales, decreasing to 0.1 for larger programs (Mayer et al. 1998). An estimated C = 0.17 has been reported for a small-scale trial of irradiated Mediterranean fruit fly (*Ceratitis capitata*), increasing to 0.42 if the males were exposed to ginger root oil (Shelly et al. 2007). The authors also observed that the competitiveness values varied inversely with the S / W ratio. In a large trial included in a successful Medfly control program, no significant induced sterility was observed until sterile/wild males ratios reached 100:1 or higher with an estimated competitiveness in the range 0.0001–0.001 (Rendón et al. 2004).

A negative correlation between the S/W males ratio and the CIS index value similar to our finding was observed in previous trials conducted under semi-field and field conditions using irradiated and transgenic sterile males (Harris et al. 2011, Damiens et al. 2016). Understanding the reasons for this negative correlation will be instrumental in the proper planning of future mosquito control programs that rely on genetic control. Actually, there is probably an optimal range of S/W males ratio, beyond which the increase in the dose of sterile males released does not add a strong benefit in the induced sterility. This is in line with the marginal decreasing productivity law and supports the empirical observation about the difficulty to achieve eradication. It is important to identify this optimal S/W ratio as this will allow accurate estimates of sterile male release densities to maximize the cost-efficiency of SIT campaigns aimed at suppression.

Finally, the mean CIS field index observed in our studies resulted in the range of the CIS values obtained in successful programs against other insect species, indicating that the SIT technology applied to *Aedes* mosquito control could achieve satisfactory results. However, the strong variability shows that the various processes of the SIT package, namely mass-rearing, handling, irradiation and release of the sterile males should be mastered to reach the highest values obtained in our study, in order to optimize cost-benefit in the field.

## Acknowledgments

We wish to thank the Municipalities of Baricella (BO), Correggio (RE), Albinea (RE), San Giorgio di Piano (BO), Crevalcore (BO) for the kind assistance in the field implementation of the trials. The field pilot trials producing the data evaluated in this study were supported by the International Atomic Energy Agency, the Emilia-Romagna Regional Bureau and the Iren S.P.A. The paper was prepared in the frame of the EU-funded research infrastructure project Infravec2-Research infrastructures for the control of insect vector borne diseases (2017 – 2021).

## Author contributions

RB, MC, FB and JB conceptualized the study. MC conducted the statistical analysis. RB, MC, FB, AP, MM and JB contributed to the writing of the original draft and to the review & editing process.

## Notes

### Competing Interest Statement

The authors have declared no competing interest.

